# Brain injury contributes to dopaminergic neurodegeneration, Lewy body pathology, and Parkinsonism preclinically with outcomes altered by T cell modulation

**DOI:** 10.1101/2025.09.16.676659

**Authors:** Colin Kelly, Julia P. Milner, Brooke A. Lester, Samantha Brindley, Lyann Hernandez, Sahitya Ranjan Biswas, Laura Murdaugh, Xiaoran Wei, Michelle L. Olsen, Matthew W. Buczynski, Alicia M. Pickrell

## Abstract

Traumatic brain injury (TBI) increases the risk of Parkinson’s disease (PD) development later in life, but much remains unknown regarding the mechanisms driving this relationship. A single, mild brain injury triggers resident and peripheral neuroinflammatory pathways that are similarly activated in PD patients, which could possibly increase susceptibility to neurodegeneration. In this study, we used preclinical mouse models of mild TBI (mTBI) and PD to evaluate how injury-induced immune signaling may exacerbate PD-associated pathologies. Dopaminergic (DA) neurons showed upregulation of genes associated with neuroinflammation, adaptive immunity, and PD following mTBI. mTBI caused degeneration of DA neurons in the substantia nigra (SN) and increased the spread of Lewy body (LB) pathology to other brain regions, such as the ipsilateral cortex, in a preclinical model of PD. Reducing adaptive immune infiltration into the central nervous system (CNS) with a transgenic model lacking mature lymphocytes, or directly by *in vivo* depletion of T cells or B cells individually, improved neurodegenerative outcomes of DA neurons following brain injury. Our results indicate the possibility of a sustained, chronic peripheral immune cell infiltration which negatively affects both DA neurons and alpha synuclein (α-syn) fibril propagation, providing insight on therapeutic windows to reduce DA neuron vulnerability.

## Introduction

Traumatic brain injury (TBI) is a highly prevalent neurological disorder [1]. While TBI is initially considered an acute injury, the secondary injury that occurs in the weeks to months following the initial insult, constituting chronic pathology, has been increasingly recognized as a contributor to prolonged symptoms and a major risk factor for further disease development. This secondary injury, characterized by the neuroinflammatory events that occur around the injury site, involves the release of cytokines and chemokines in addition to a complex crosstalk between resident and infiltrating peripheral immune cells that play both protective and neurotoxic roles [2].The neuroinflammatory events that occur and persist well after TBI share many similarities to the neuroinflammatory events that occur in neurodegenerative diseases, such as Parkinson’s disease (PD).

PD is the most common motor deteriorating neurodegenerative disease, resulting in 90,000 diagnoses each year in the United States with over 10 million current diagnoses worldwide [3]. Patients are clinically diagnosed with PD when they exhibit cardinal motor signs and symptoms of muscle rigidity, bradykinesia, postural instability, and tremors, which are attributed to the progressive degeneration of dopaminergic (DA) neurons within the substantia nigra (SN) [4]. Another pathological identifier of the disease is the cytoplasmic neuronal inclusions, termed Lewy Bodies (LB) and Lewy Neurites, which form from abnormal deposits of alpha synuclein (α-syn) protein [5]. α-syn has a propensity to misfold and aggregate into insoluble fibrils, spread cell-to-cell, and recruit endogenous monomeric α-syn for increased propagation and aggregation [6].

While aging remains the greatest risk factor for PD development, there is growing interest in other genetic and environmental factors that contribute to risk, such as TBI [7]. The occurrence of TBI positively correlates with development of PD later in life – one study following over 300,000 veterans showed that mild TBI (mTBI) increased PD diagnosis likelihood by 56% [8]. Another more recent study confirmed this association between injury and PD development [9]. However, the causal cellular and molecular mechanisms that link mTBI and PD are not yet understood.

It is well established that patients with PD have increased levels of proinflammatory cytokines and altered levels of circulating immune cells in both their brain and serum [10–13]. α-syn aggregates have been shown to be a driving force in this immune activation, further advancing disease pathology [5]. This makes the neuroinflammatory cascade triggered by mTBI particularly interesting regarding the potential influence it may play on DA neuron vulnerability and the progression of PD-associated LB histopathology. Few studies have adequately addressed whether the secondary injury associated with mTBI contributes to the pathophysiological changes associated with PD or other neurodegenerative diseases, especially when evaluating chronic time points post-injury. In more recent studies, the role of the adaptive immune system, specifically infiltrating T cell subsets, has been studied in the context of PD. Data from preclinical models of PD show that CD4^+^, CD8^+^, and even gamma delta (γδ) T cell populations enter the brain and play a role in mediating neuroinflammatory pathways that negatively impact DA neuron degeneration [14–16]. *In vitro*, catecholaminergic murine primary neurons have been shown to be specifically susceptible to T cell mediated degeneration through the induction of major histocompatibility complex class I (MHC-I) via interferon gamma (IFN-γ), which is a cytokine released by adaptive immune cells during the neuroinflammatory response [17]. How adaptive immune cell populations, which have been shown to persist in the central nervous system (CNS) at chronic time points post-TBI [18], impact the pathophysiological changes in PD in a rodent model of mTBI has yet to be determined.

To address these questions, we utilized a single hit, diffuse, closed head injury paradigm, both separately and in conjunction with a preclinical model of PD, to assess chronic functional outcomes to DA neuron health and behavioral consequences. Previous studies have not examined the effect of a prior, single-hit mTBI on DA degeneration and α-syn propagation in a model of rodent PD. We further evaluated the potential role of adaptive immune signaling by applying these same approaches with immunodeficient Rag2 KO animals and animals with antibody depleted T and B cells to uncover the contribution that adaptative immunity participates in DA neurodegeneration in the contexts of injury. Our findings contribute to a strengthened understanding concerning the potential lymphocyte-driven degeneration of DA neurons, uncover the contribution of neuroinflammation-induced injury to susceptible neuronal populations, and provide insight into how mTBI affects α-syn accumulation and propagation.

## Results

### Closed-head diffuse, single hit mTBI results in Parkinsonian-like degeneration of dopaminergic neurons

Recent studies have demonstrated that rodent TBI can sufficiently cause loss of DA populations in the SN [19]. However, many TBI models do not appropriately reflect the injury severity typically seen in cases of military veterans, sports collisions, or civilian accidents. While cortical contusion [20, 21] and lateral fluid percussion [22] models produce TBIs that allow researchers to damage the intended brain side directly, these craniotomy-based approaches range from moderate to severe and are oftentimes penetrating, requiring removing the skull to produce an open head injury that is less comparable to the closed head mTBIs more commonly experienced by military and civilian populations. Moreover, impact injury models, such as weight drop injury, require multiple hits that occur over a short time frame (hours/days) to produce sufficient brain damage [23]. This multiple-impact approach is poorly reflective of the most commonly experienced brain injuries, where individuals predominately experience a single impact injury that damages the brain.

To address the challenges posed by previous TBI studies that have evaluated PD like pathologies, we implemented a brain injury mouse model in our laboratory that directly impacts areas of the brain implicated in common head injuries [24]. Here, we generated a single, mild closed-skull impact injury aimed at an area of skull above the prefrontal cortex and striatum (STr). The parameters of this hit result in low mortality (1.5% over 100+ surgeries) and an absence of skull fractures (see Material and Methods). Our injury conditions resulted in a tail flinch at the time of impact, coupled with minor bruising under the skull, but no lesions or major hemorrhage.

We first aimed to determine whether our mTBI model had negative consequences on DA neuronal health. We evaluated the number of DA neurons in the SN 90 days post-injury (dpi) and found a significant, 30% decrease in the number of tyrosine hydroxylase (TH^+^, a protein marker of DA biosynthesis) neurons following mTBI (Fig. 1A,B). These neurons have axonal projections that travel from the SN to the STr where dopamine is released, so we measured the optical density of these fibers in the STr, and, consistent with the loss of DA neurons in the SN, found a decrease in TH^+^ fluorescent intensity (Fig. 1C,D). To verify that these observed changes to DA neurons are degenerative as opposed to a downregulation of TH, total cells (DAPI^+^) were counted within the SN region displaying a similar, significant reduction (Fig. 1E). Considering the alterations found in the STr, we examined whether injured animals showed loss of striatal neurons, or if our observations were specific to DA neurons. Nissl^+^ neurons were quantified from the STr, a region predominantly characterized by GABAergic neurons, which showed no significant differences between injured animals and shams (Sup. Figure 1A).

**Figure 1.**
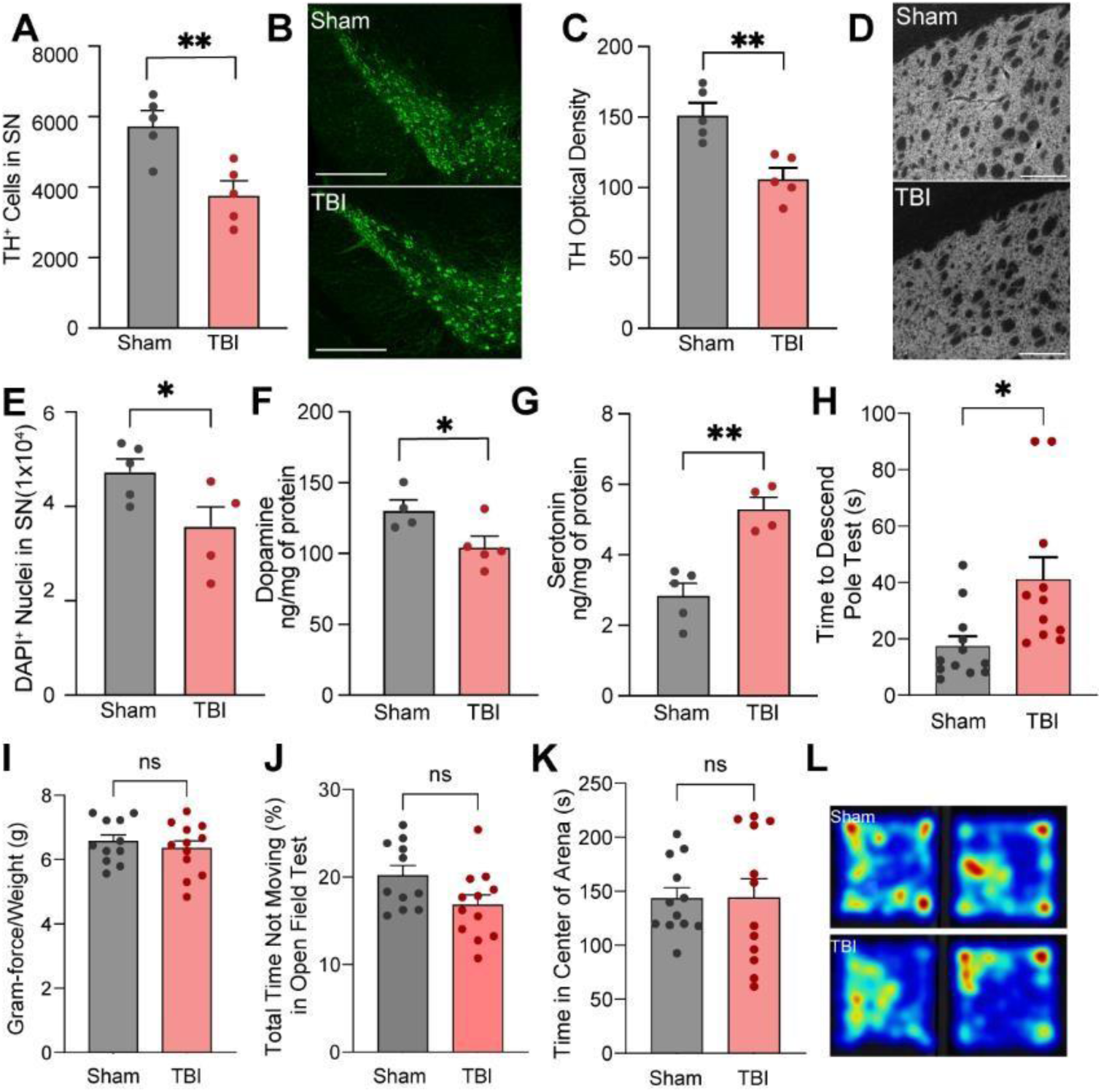
SN DA neurons degenerate following mTBI. *A*. Quantification of TH^+^ neurons within the SN 90 d after mTBI. *B*. Representative confocal images of sham and TBI SN 90 d after injury. Tissue was stained for TH (green) to indicate DA neurons. Scale bar = 400µm. *C*. Quantification of TH^+^ optical density in STr 90 d after injury. *D*. Grayscale representative confocal images of sham and TBI STr 90 d after injury. Tissue was stained with TH to indicate DA neuron fibers. Scale bar = 200µm. *E*. Quantification of DAPI^+^ nuclei within the SN of sham and TBI mice 90 d post-injury. *F*-*G*. HPLC analysis of neurotransmitters dopamine and serotonin in STr tissue of sham and TBI mice 90 d after injury. *H*. Pole test performance of sham and TBI animals 90 d after injury. *I*. Grip test performance of sham and TBI animals 90 d after injury. *J-K*. OFT performance of sham and TBI animals. Time not moving (*J*) (% of total time) and time in the center (K) of the arena were evaluated 90 d after injury. *L*. Representative heatmaps of open field activity. Each dot represents one animal. \**p* < 0.05; \*\**p* < 0.01; \*\*\**p* < 0.001; analyzed by unpaired t-test. ns = not significant. Error bars = ± SEM.

**Supplementary Figure 1.**
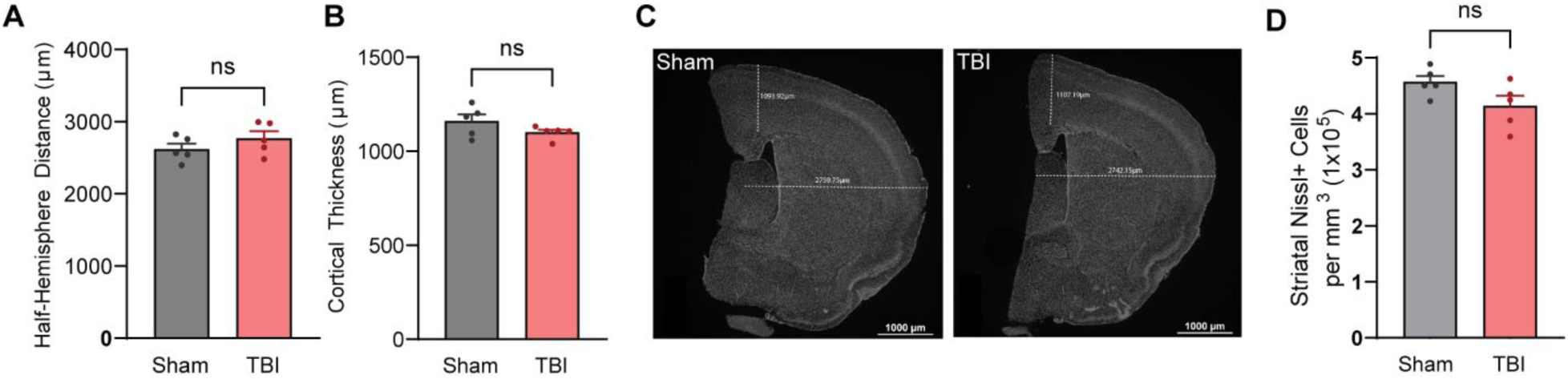
Cortical thickness and brain size are unaffected following TBI at 90 days. *A.* Quantification of Nissl^+^ neurons in the STr 90 dpi. *B*. Quantification of hemisphere distance, measured as a horizontal line from the lateral ventricle to the cortex. *C*. Quantification of cortical thickness, measured as a vertical line from the lateral ventricle to the cortex. *D*. Representative images of sham and mTBI brains used for analyses. n = 5 for each group. Each dot represents one animal. \**p* < 0.05; \*\**p* < 0.01; \*\*\**p* < 0.001; analyzed by unpaired t-test. ns = not significant. Error bars = ± SEM.

Next, we evaluated whether the loss of DA neurons caused alterations in neurotransmitter concentrations within the STr. mTBI animals showed a significant decrease in dopamine concentration compared to sham controls (Fig. 1F). Interestingly, these same animals showed an increase in serotonin levels (Fig. 1G), which may be a compensatory mechanism consistent with other preclinical PD rodent models [25]. To test whether the deficits in DA neuron loss impacted motor function, animals underwent a series of behavioral assessments. Injured animals demonstrated an increased latency to complete the pole test, though there were no differences on the grip strength exam, supporting a deficit in motor coordination unrelated to overall strength (Fig. 1H,I). At this 90 days (d) time point, these same groups performed open field testing, where there were no significant differences in either their time spent not moving or their time spent in the center of the arena (Fig. 1J-L).

We sought to determine whether our observed loss of DA neurons was indeed a degenerative process and not an immediate result of the initial insult. To do this, we examined DA neuron survival 14 dpi. Sham and mTBI animals displayed no significant DA cell death at this subacute timepoint, additionally supported by no differences in DAPI^+^ cells within the same contour of the SN (Fig. 2A-C). Likewise, when analyzing the STr of these animals, we found no differences in the optical density of DA fibers (Fig. 2D,E). Nissl^+^ neurons in the STr were not significantly different between injured and sham animals (Fig. 2F). We also assessed whether our injury resulted in any brain deformation and found no significant alterations in brain morphology, cortical thickness, and hemisphere width 14 dpi (Sup. Figure 1B-D).

**Figure 2.**
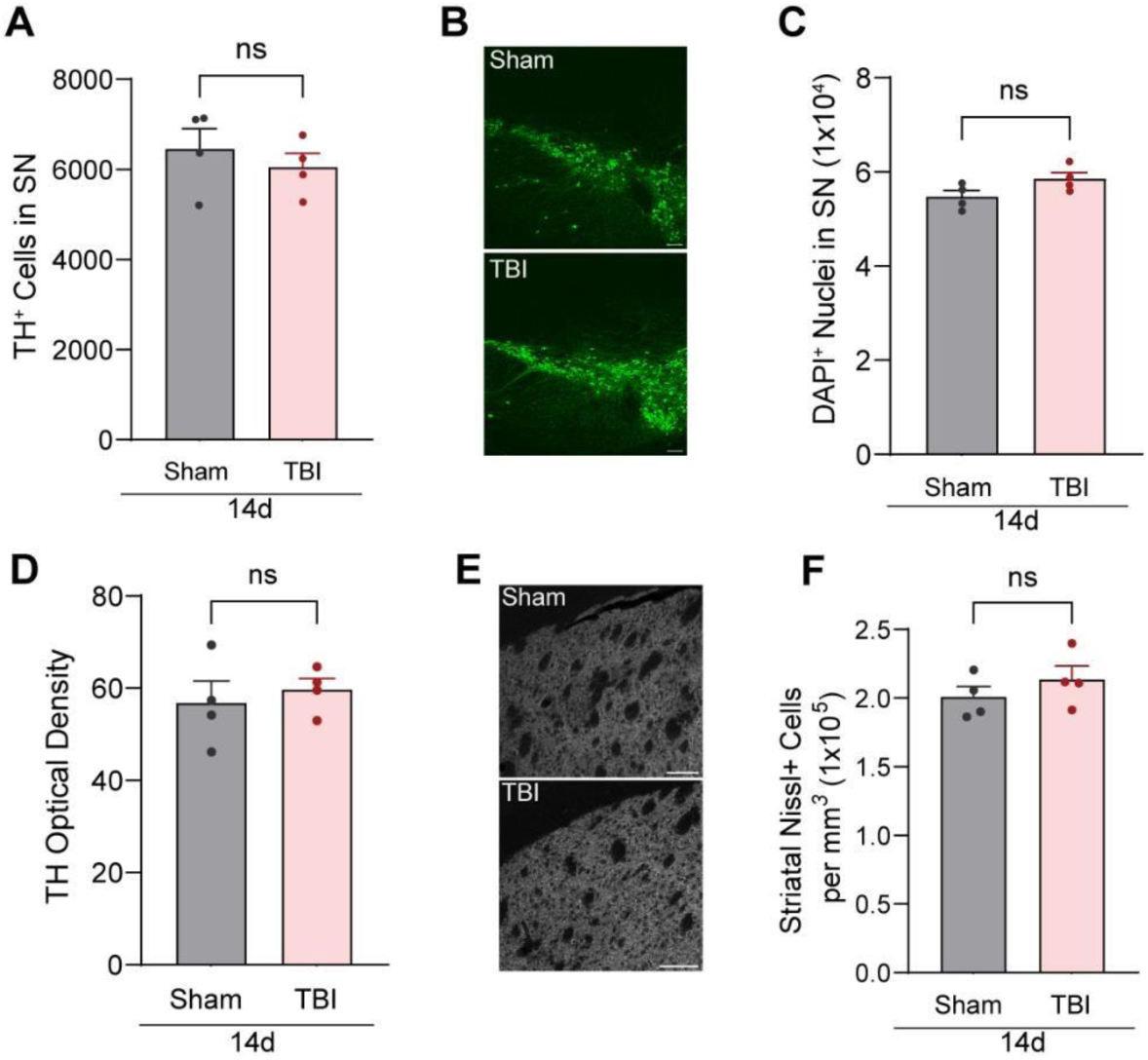
mTBI does not result in DA neurodegeneration at a subacute time point postinjury. *A*. Quantification of SN TH^+^ neurons 14 d after TBI. *B*. Representative confocal images of 14 d sham and mTBI SN. Tissue was stained for TH (green) to indicate DA neurons. Scale bar = 100µm. *C*. Quantification of DAPI^+^ nuclei within the SN 14 dpi. *D*. Quantification of STr optical density 14 d after injury. *E*. Grayscale representative confocal images of sham and mTBI STr 14 d after injury. Tissue was stained for TH to indicate DA neuron fibers. Scale bar = 100µm. *F*. Quantification of Nissl+ cells in the STr 14 d after injury. n = 4 for each group. Each dot represents one animal. \**p* < 0.05; \*\**p* < 0.01; \*\*\**p* < 0.001; analyzed by unpaired t-test. ns = not significant. Error bars = ± SEM.

Lastly, to evaluate the scale of immune response to our injury, transcriptomic profiling of cortical tissue was performed 2 hours (h) after injury and revealed a distinct clustering between samples that included a significant upregulation of immune activation and immune signaling signatures in injured mice as compared to shams (Sup. Figure 2A). Using a p < 0.05 and a threshold of 1.5 fold-change as a cutoff, our analysis identified 446 differentially expressed genes (DEGs), with most of these genes being upregulated and playing a role in the early immune response (Sup. Figure 2B, C, Table S1). We next compared our upregulated gene set to the upregulated DEGs found in our previous data that employed a more moderate model of controlled cortical impact (CCI) brain injury requiring a craniotomy [26]. While moderate CCI resulted in a greater number of upregulated DEGs, we found a large overlap of 71 upregulated genes between models, suggesting a successful, albeit less severe neuroinflammatory response with our current paradigm (Sup. Figure 2D). Taken together, these data indicate that our model of mTBI results in a truly mild injury that induces DA degeneration chronically post-injury.

**Supplementary Figure 2.**
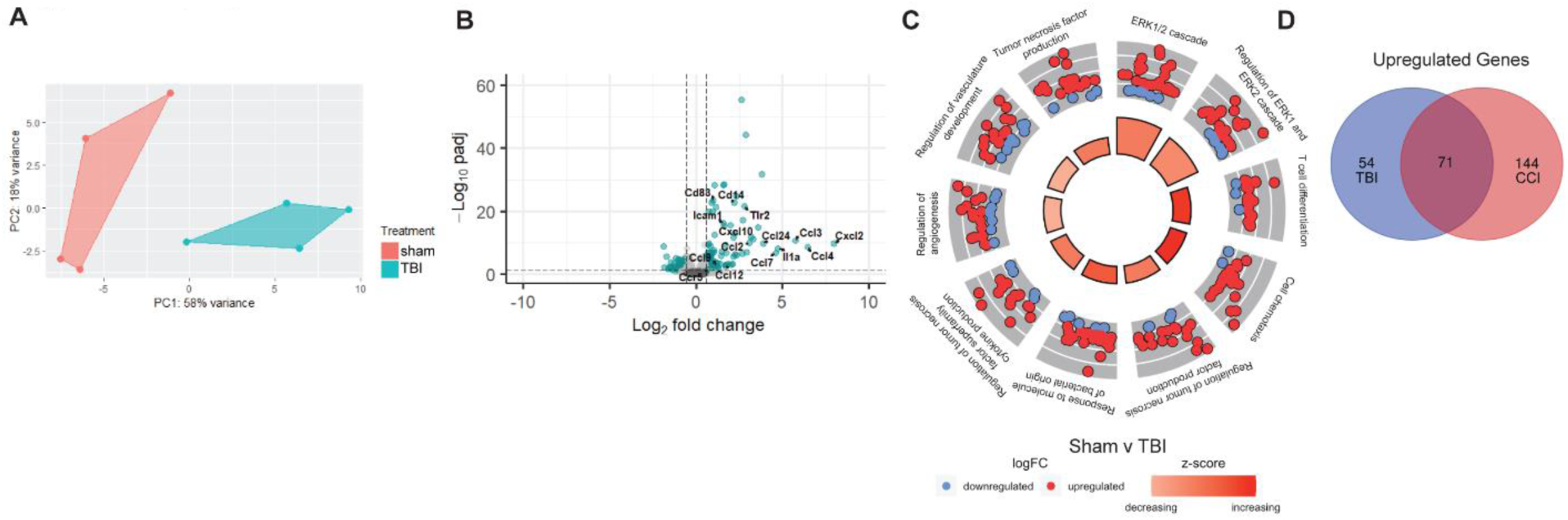
Transcriptomic profile of cortical tissue reveals significant upregulation of peripheral immune cell signaling signatures 2 h after injury. *A*. Principal component analysis (PCA) plot of samples used in RNAseq analysis. n = 4 for sham and 4 for TBI. *B*. Volcano plot of significant DEGs. *C*. Genes from injured cortex revealed GO terms associated with chemotaxis, T cell migration, and ERK1/2 signaling. *D*. Venn diagram showing overlap of significant, upregulated genes in TBI compared to genes altered 2 hours after cortical contusion (CCI) injury.

### mTBI increases severity of α-syn propagation in a preclinical model of PD

Considering that mTBI alone results in an apparent chronic neuroinflammatory process that negatively impacts DA neurons, we next wanted to test whether eliciting mTBI prior to seeding a preclinical model of rodent PD accelerated associated pathologies. Because DA neurons are considered susceptible to adaptive immune-related signaling mechanisms, and neuroinflammation occurs both in the secondary injury following mTBI and in the progression of PD, we would expect a pre-existing neuroinflammatory event to play a detrimental role in the progression of disease pathology. To test this, we created a combinational model using our mTBI paradigm coupled with the α-syn preformed fibril (PFF) model of PD. Double treatment animals received mTBI 30 d prior to intrastriatal injection of PFFs and were sacrificed 60 d post-injection. 30 d allows enough time for the activation, infiltration, and downstream effects of both the resident and peripheral immune response post-injury, prior to the seeding of PFFs.

We first validated successful injection site seeding of α-syn fibrils in the STr (Fig. 3A). From here, fibrils are retrogradely transported via axonal projections to the SN, where they continue aggregation and propagation in DA neurons causing degenerative cell death. Using serial sectioned midbrain tissue, we visualized and quantified DA neurons in the SN co-labeled with phosphorylated-α-synS129 (p-α-syn) to detect the formation of aggregates/LBs (Fig. 3B). Quantification of DA neurons revealed a significant decrease in the mTBI + PFF group compared to shams or PFF only groups, with no difference in DA populations between sham and PFF animals, which was further validated by quantification of DAPI^+^ cells (Fig. 3C,D). Additionally, mTBI + PFF animals had a significantly lower percentage of DA neurons that were co-labeled with p-α-syn than mice that had only received the PFF treatment (Fig. 3E). We also quantified the number of TH^-^ DAPI^+^ cells within the SN, considered non-DA cells, that were p-α-syn^+^, finding no difference between the PFF only and mTBI + PFF groups (Fig. 3F). However, we did note that significant cortical spreading of p-α-syn^+^ cells occurred in injured mice as opposed to those only exposed to PFFs, indicating that brain injury did exacerbate LB pathology (Fig. 3G,H). Optical density analysis from STr tissue stained with TH showed reduced fiber fluorescence in the double treatment group, whereas PFF only animals showed no significant decrease compared to controls (Fig. 3I,J). We then performed high-performance liquid chromatography (HPLC) analysis on striatal tissue from the double treatment group and controls, finding a consequential decrease in dopamine concentration that aligned with the loss of DA neurons in the SN and reduced intensity of DA fibers in the STr for mTBI + PFF mice (Fig. 3K). These data suggest that mTBI prior to PFF seeding contributes to increased propagation of α-syn fibrils, but that at this time point (60 d post-injection), DA neuron death appears to be mTBI driven, as both the mTBI only group and the mTBI + PFF group show between ∼30-35% cell death.

**Figure 3.**
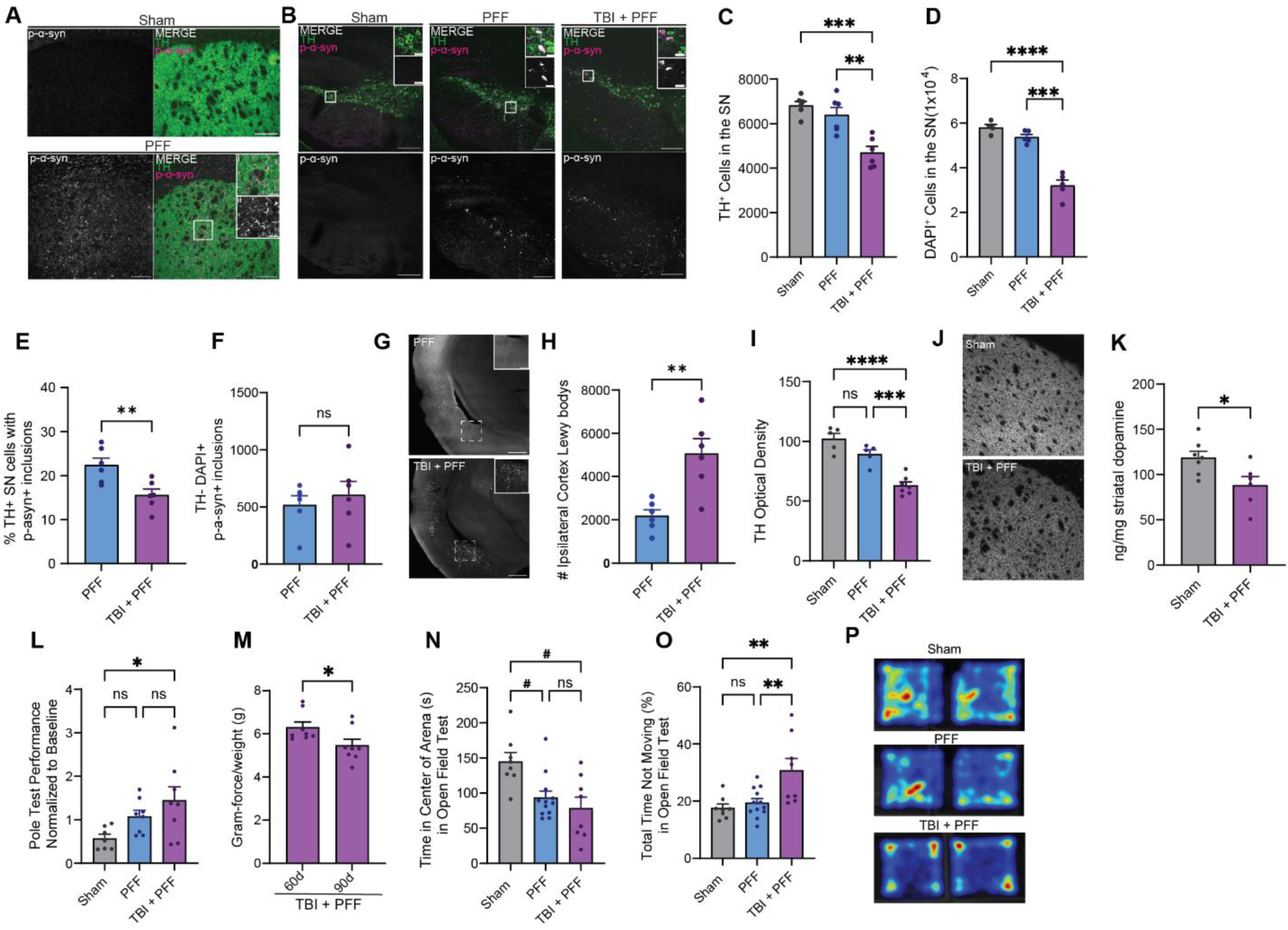
mTBI alters α-syn pathology. *A*. Representative confocal images of sham and PFF injected STr 60 days post-injection. Tissue was stained for TH (green) to indicate DA fibers and p-α-syn129 (magenta) to indicate α-syn fibrils/inclusions. Scale bar = 200µm. Inset scale bar = 50µm. *B*. Representative confocal images of SN in sham, PFF, or TBI + PFF animals. Tissue was stained for TH (green) to indicate DA neurons and p-α-syn129 (magenta) to indicate α-syn fibrils/inclusions. Scale bar = 200µm. Inset scale bar = 20µm. *C.* Quantification of SN TH^+^ neurons in sham, PFF, and TBI + PFF treated animals. *D.* Quantification of DAPI^+^ nuclei within the SN in sham, PFF, and TBI + PFF treated animals. *E.* Quantification of the percentage of DA neurons with p-α-syn129 inclusions and *F.* the number of non-DA neurons with p-α-synS129 inclusions within the SN in PFF and TBI + PFF treated animals. G. Representative confocal images of LBs in the ipsilateral cortex stained with p-α-syn129. Scale bar = 200µm. *H.* Quantification of LBs in the ipsilateral cortex. *I*. Quantification of optical density of TH in the striatum. *J.* Representative confocal images of striatum tissue stained with TH. *K.* HPLC analysis of striatal dopamine concentration in sham and TBI + PFF treated animals. *L*. Pole test performance of sham, PFF, and TBI + PFF animals 90 d after initial surgery. *M*. Comparison of grip strength for TBI + PFF animals at 60 and 90 d post-initial injury. *N-P*. OFT measuring (*N*) total time in the center of the arena, (*O*) percentage of total time spent not moving, and (*P*) a heatmap representation of animal movement and location. n = 6 for quantification data, n = 8-12 for behavioral assessments. Each dot represents one animal. *^,#^*p* < 0.05; **^,##^*p* < 0.01; ***,^###^*p* < 0.001; analyzed by one-way ANOVA, unpaired students t-test, and the Kruskal-Wallis test. ns = not significant. Error bars = ± SEM.

To determine if these findings culminated in reduced motor performance or other behavioral deficits, animals performed the pole test, grip strength test, and open field test. mTBI + PFF mice showed reduced capabilities on the pole test, indicative of reduced motor coordination (Fig. 3L). While the grip strength test showed no significant differences between experimental groups, mTBI + PFF animals were the only group that had reduced in-group performance between the 60 d and 90 d time points (Fig. 3M). Open field test revealed that mTBI + PFF animals spent significantly less time in the center of the arena than sham animals, but there was no significant difference when compared to the PFF only group. However, the mTBI + PFF animals spent significantly more time not moving than both other groups (Fig. 3N,O,P), indicating anxiety-like behavior – a behavior that was absent in mice that received mTBI alone.

### mTBI causes transcriptional changes associated with peripheral immune signaling and PD in DA neurons

To gain better insight into how DA neurons responded to mTBI, we assessed transcriptomic gene expression changes after injury. Because single nuclei RNA sequencing and bulk RNA sequencing methods will overlook changes occurring to DA neurons due to their complex morphology and long STr projections, we employed the use of RiboTag mice that have an HA tag on the 60S ribosomal subunit controlled by a floxed STOP codon. Crossing the RiboTag line with a dopamine transporter (DAT) cre line removes the STOP codon to specifically allow for the direct pulldown of tagged polysomes from DA neurons [27] (Fig. 4A). STr tissue was collected from sham or mTBI RiboDAT animals 90 dpi and subsequently processed by immunoprecipitation. We confirmed that IPs harvested from DA neuron populations were enriched by analyzing mRNA expression of DAT and TH via qPCR of IP versus whole brain control samples (Fig. 4B). With the RNA collected from IPs, we performed a whole transcriptome RNA-seq expression analysis to evaluate what transcriptomic alterations, if any, occur following mTBI within these cell populations that might elucidate the processes or pathways driving the degeneration. RNA-seq analysis revealed 291 DEGs using p < 0.05 and a threshold of 1.2 fold-change in expression as a cutoff for analysis (Fig. 4C, Table S2). Interestingly, many of the top significantly downregulated genes in injured animals were directly related to DA survivability and health (*Th*, *Fgf20*, *Snai1*, *En1*, *Chrm5*). *Fgf20,* which showed a 25-fold decrease in mTBI animals, is preferentially expressed in the SN and has been shown to significantly protect against the loss of DA neurons in a rat model of PD [28]. One of the most significantly upregulated genes following injury, *Trib3*, is highly elevated in DA SN neurons of PD patients and has been shown to play a role in mediating cell death and neurodegeneration in preclinical PD models [29]. The induction of *Trib3* has been observed prior to measurable loss of DA neurons in these PD models. *Th* was downregulated following mTBI, which was validated in our immunohistochemistry experiments where mTBI mice showed decreased TH^+^ cell counts and reduced optical density.

**Figure 4.**
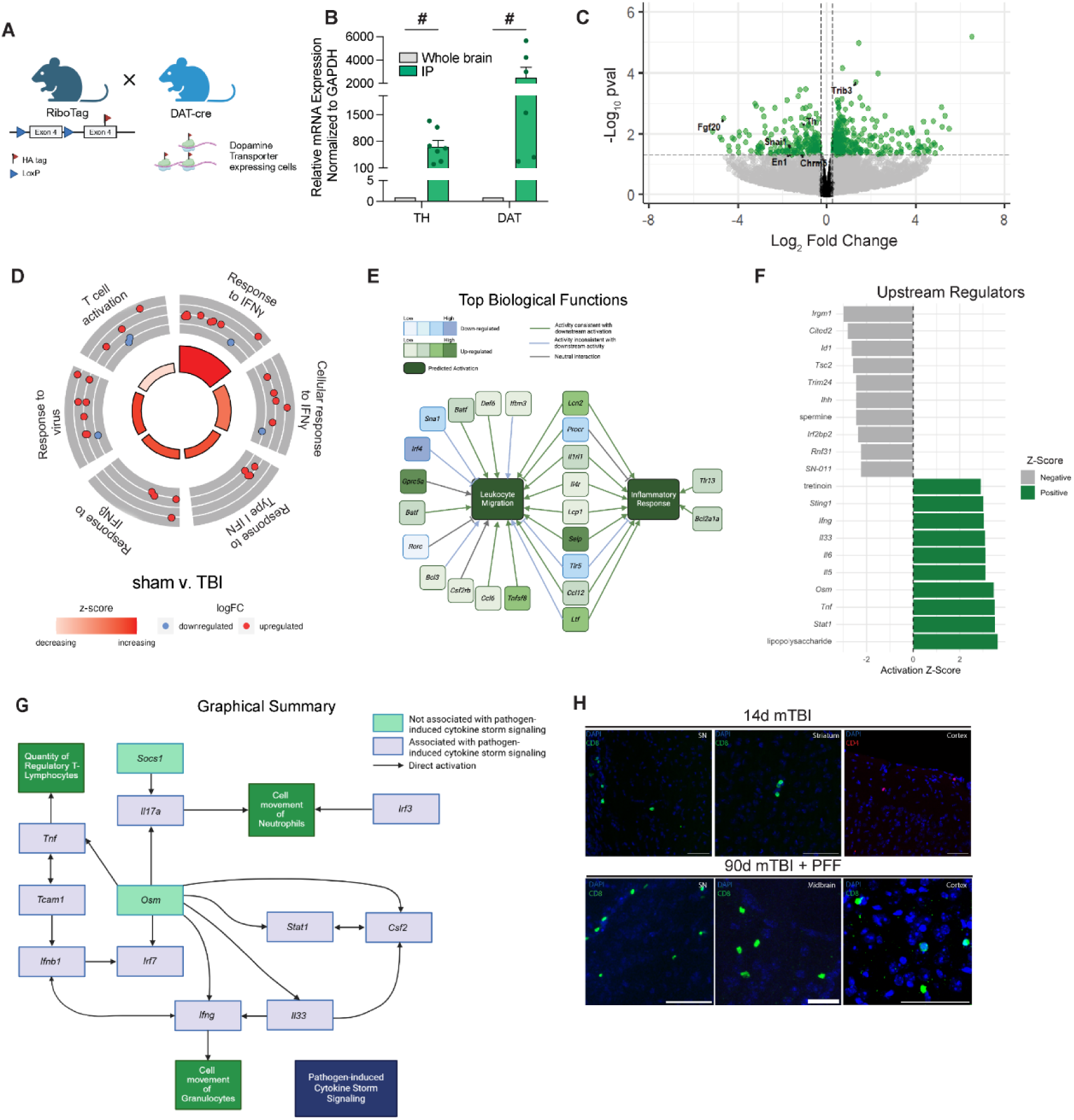
Transcriptomic changes from DA neurons are associated with PD, neuroinflammation, and peripheral immune signaling 90 d following injury. *A*. Cartoon diagram for the generation of RiboDat animals. Panel made with BioRender. *B*. qPCR of mRNA expression for TH and DAT normalized to GAPDH after immunoprecipitation of HA^+^ ribosomes compared to whole brain RNA from the same animal. n = 6-7 animals. *C*. Volcano plot of significant upregulated and downregulated genes in mTBI animals 90 d after injury. n = 4 TBI and 3 sham. *D*. GO terms associated with T cell activation and response to IFN-γ. *E*. Pathway analysis of the top biological functions related to the DEG list. *F*. Top 10 upregulated and downregulated upstream regulators of DEGs. *G*. Graphical summary of the pathway most implicated in relation to DEGs. *H*. Representative confocal images of 14 d mTBI and 90 d mTBI + PFF tissue. Tissue was stained for TH (DA neurons), CD8 (CD8 T cells) and CD4 (CD4 T cells). Each dot represents one animal. ^#^*p* < 0.05; ^##^*p* < 0.01; ^###^*p* < 0.001; analyzed by one-sample Wilcoxon signed-rank test. ns = not significant. Error bars = ± SEM. Scale bars = 50µm and 100µm.

Further analysis of the expression profile of these DEGs using gene ontology (GO) revealed a significant increase in the expression of genes related to T cell activation and response to IFN-γ, implicative of the possible role of the adaptive immune system (Fig. 4D). Further examination using Ingenuity Pathway Analysis (IPA) showed that top biological functions of our dataset were leukocyte migration and inflammatory response, supporting the potential role of adaptive immunity in DA neuron susceptibility within these processes (Fig. 4E). We examined upstream regulators of DEGs, and found that both the top upregulated hits (*Sting1*, *Ifng*, *Tnf*, *Il6*, LPS, etc) and the top downregulated (*Irgm1*, *Cited2*, *Trim24*, etc) are strongly associated with neuroinflammatory pathways that can play a detrimental role in DA neuron susceptibility (Fig. 4F). Additionally, the graphical summary generated with this data, with its overarching emphasis being pathogen-induced cytokine storm signaling, further indicated a chronic neuroinflammatory activation (Fig. 4G).

To further examine the findings supporting that adaptive immunity may be involved in the observed DA susceptibility, we performed immunohistochemistry (IHC) on both 14 d mTBI and 90 d mTBI + PFF brain tissue for CD8, CD4, and CD20 antibodies to examine whether T and B cells infiltrated and remained in the brain parenchyma. While we never detected a significant infiltration of B cells, there was an initial infiltration of both CD4^+^ and CD8^+^ T cells at 14 d in both forebrain and midbrain tissue (Fig. 4H). 90 d mTBI + PFF tissue showed a consistent presence of CD8^+^, but not CD4^+^ T cells around the SN and cortex (Fig. 4H). Taken together, these data support a chronic process of degeneration following injury that appears to be mediated by the role of secondary neuroinflammatory injury mechanisms driven by adaptive immunity.

### Adaptive immunodeficiency is neuroprotective against the effects of mTBI + PFF treatment

Considering that transcriptomics of injured animals revealed interactions with interferon-γ and implicated the adaptive immune response, we next decided to test our injury paradigm with immunodeficient mice. To do this, we used Rag2 KO mice deficient in both mature T and B cells. We found that mTBI + PFF Rag2 KO animals showed significant protection against declines in motor coordination on the pole test compared to the deficit seen in WT animals (Fig. 5A). To confirm that the decline in ability of wildtype (WT) mice was truly dopamine dependent, we injected the same cohort with a levodopa (L-DOPA) cocktail and found that they significantly recovered function in the task (Fig. 5B). Following sacrifice, DA populations in the SN were quantified. We found that Rag2 KO animals were protected from the cell death observed in WT animals, further confirmed by quantification of DAPI^+^ nuclei (Fig. 5C,D). While immunodeficient animals proved resistant to the degeneration of DA neurons, the propagation and seeding of α-syn fibrils was more severe. DA neurons in the SN of Rag2 KO animals showed a greater percentage of co-labeling with α-syn fibrils than those in WT mice (Fig. 5E,F). We also observed a significant increase of non-DA cells in the SN with α-syn aggregation, possibly indicating a more severe spread of α-syn fibrils (Fig. 5G).

**Figure 5.**
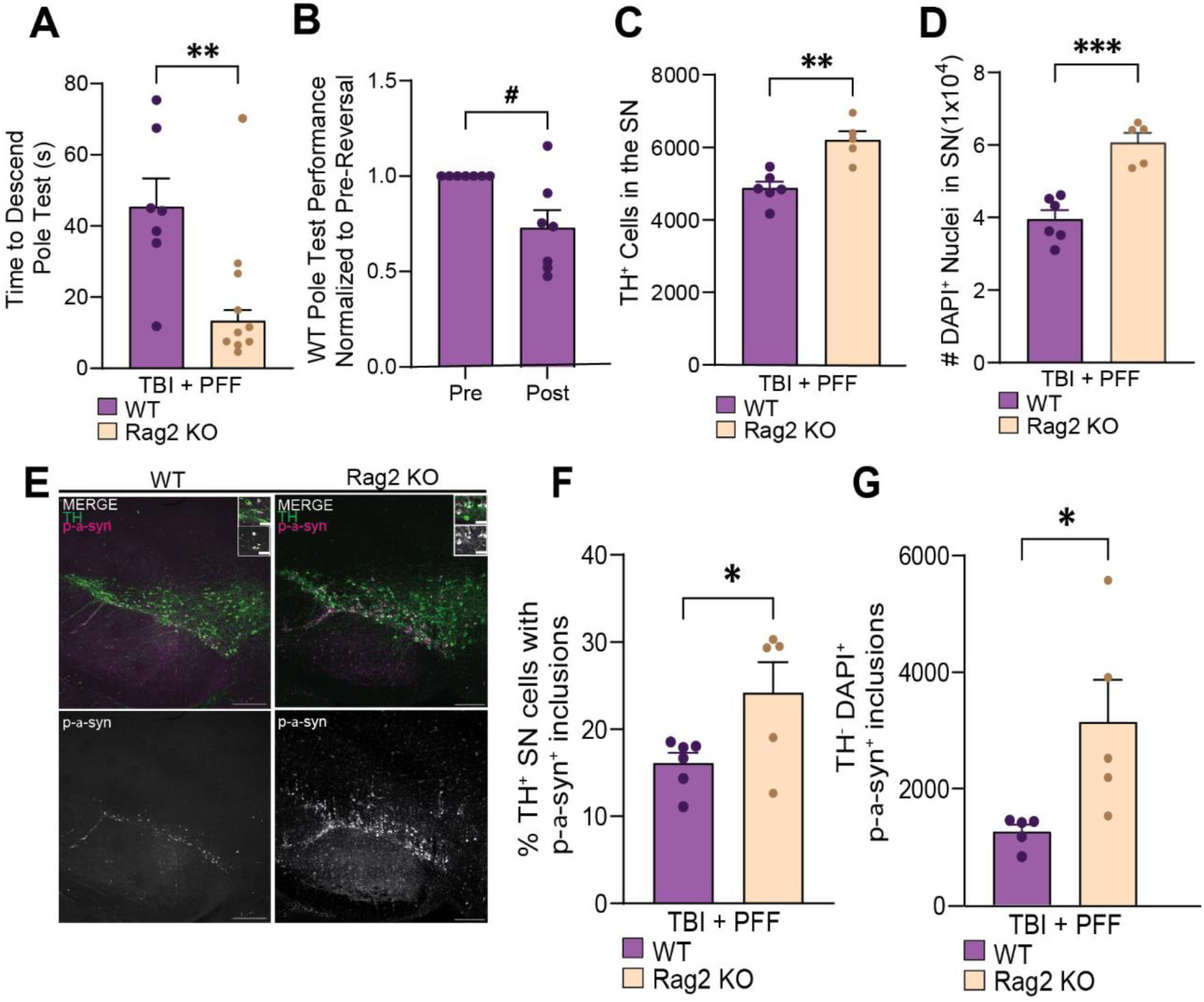
Genetic lack of mature T and B cells rescues motor deficits and ameliorates DA neuron death. *A*. Pole test performance of mTBI + PFF WT and Rag2 KO animals 90 d postinjury. *B*. Pole test performance of WT animals after L-DOPA reversal. *C*. Quantification of TH^+^ neurons in the SN of mTBI + PFF WTs and Rag2 KO animals. *D*. Quantification of DAPI^+^ nuclei within the SN of mTBI + PFF WTs and Rag2 KOs. *E*. Representative confocal images of SN in mTBI + PFF WT and Rag2 KOs. Tissue was stained for TH (green) to indicate DA neurons and p-α-syn129 (magenta) to indicate α-syn fibrils/inclusions. Scale bar = 200µm. Inset scale bar = 20µm. *F-G*. Quantification of (*F*) the percentage of DA neurons with p-α-syn129 inclusions and (*G*) the number of non-DA neurons with p-α-syn129 inclusions within the SN. n = 5-6 for quantification data. Each dot represents one animal. *,^#^*p* < 0.05; **^,##^*p* < 0.01; ***^,###^*p* < 0.001; analyzed by unpaired t-test or one-sample Wilcoxon signed-rank test. ns = not significant. Error bars = ± SEM.

Because Rag2 KO mice are deficient in both T and B cell populations from birth, we next wanted to determine whether T or B cells specifically were responsible for the susceptibility of DA neurons in our model of mTBI + PD. WT animals received mTBI, and 14 d later were split into four different groups (Fig. 6A). We chose 14 dpi as it is when we previously observed adaptive immune cell infiltration (Fig. 4H), in congruency with other groups that have reported adaptive immune cell infiltration in other injury models. Group

- received retro-orbital injections of *in vivo* CD20 antibody to deplete B cells, while Group
- received an IgG isotype matched control. These injections were done once every 30 d. Group 3 received IP injections of *in vivo* CD4/CD8 antibodies to deplete T cells, while Group 4 received an IgG isotype matched control. These injections were done once every 7 d. Controls were selected based on previous studies that used similar pharmacological depletion strategies [18, 30]. All injection protocols began 14 d after mTBI, as our data showed infiltration of adaptive immune cells at this time point while simultaneously preceding any significant DA degeneration.

**Figure 6.**
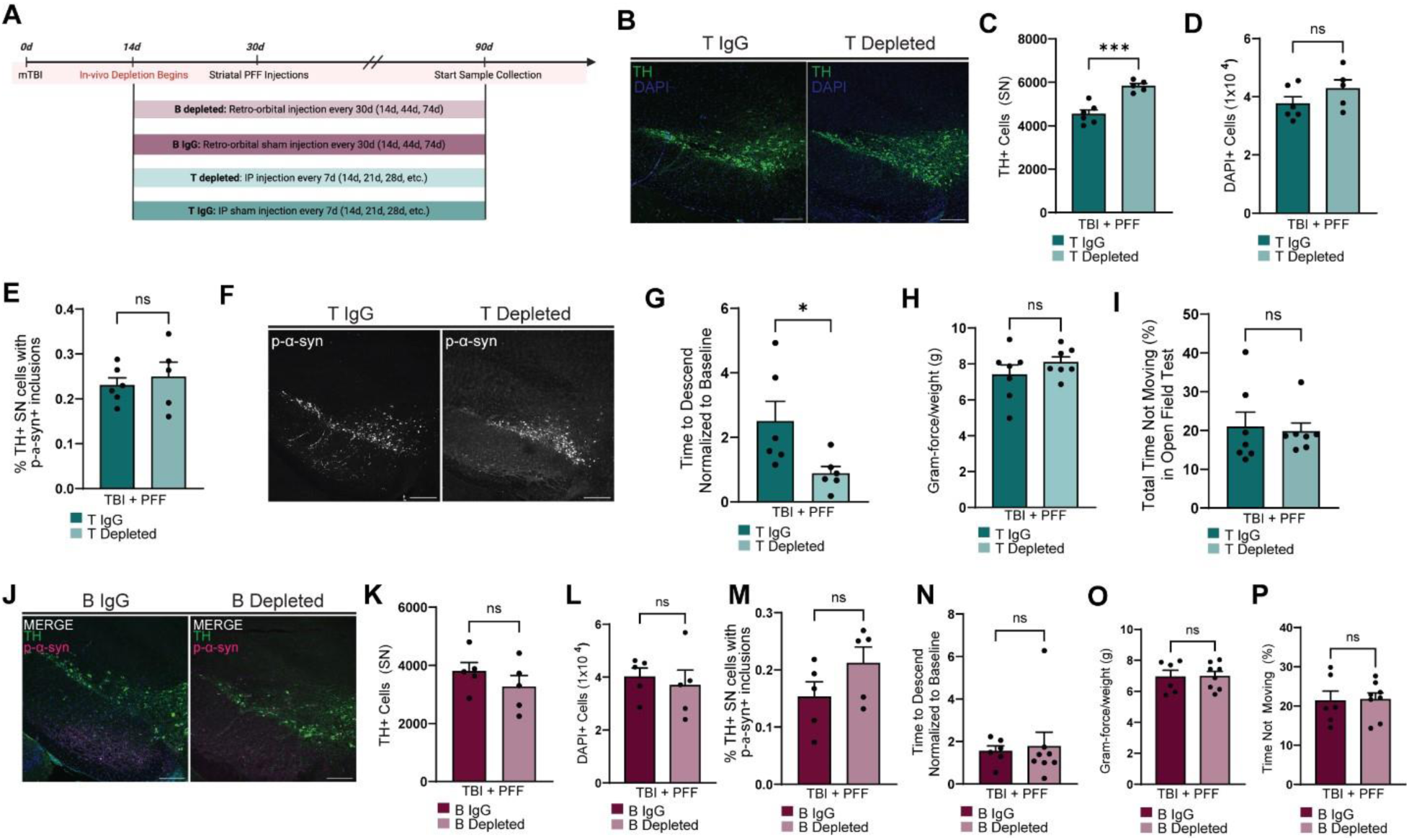
T cell but not B cell depletion 14 days post-injury reverts injury induced DA phenotypes. *A*. Schematic timeline of *in vivo* T and B cell depletion cohorts. Figure created in BioRender. *B.* Representative confocal images of T IgG and T depleted SN. Tissue was stained for TH (green) to indicate DA neurons. Scale bars = 200µm. *C*. Quantification of TH^+^ neurons in the SN of T IgG and T depleted mTBI + PFF mice. *D*. Quantification of DAPI^+^ nuclei within the SN of T IgG and T depleted mTBI + PFF mice. *E.* Quantification of the percentage of DA neurons with p-α-syn129 inclusions. *F.* Representative confocal images of T IgG and T depleted p-α-synS129 inclusions in the SN. Scale bars = 200µm. *G.* Pole test performance 90 d after initial surgery. *H*. Comparison of grip strength 90 d post-initial injury. *I*. Open field test measuring total time not moving (s). *J*. Representative confocal images of DA neurons in B IgG and B depleted SN. Scale bars = 200µm. *K*. Quantification of TH^+^ neurons in the SN of B IgG and B depleted mice. *L*. Quantification of DAPI^+^ nuclei within the SN of B IgG and B depleted mice. *M*. Quantification of the percentage of DA neurons with p-α-syn129 inclusions in the SN. *N*. Pole test performance 90 d after initial surgery. *O*. Comparison of grip strength 90 d post-initial injury. *P*. Open field test measuring total time not moving (s). n = 5-8 animals per group. Each dot represents one animal. \**p* < 0.05; \*\**p* < 0.01; \*\*\**p* < 0.001; analyzed by unpaired t-test. ns = not significant. Error bars = ± SEM.

To ensure that mice receiving either T or B cell-depleting antibodies showed a proper and sustained reduction in their respective cell population, whole blood was collected via cardiac puncture at endpoint, and samples were assessed by flow cytometry. B cell depleted mice showed a 99% reduction in total B cells compared to IgG controls (Sup. Figure 3A, B). Similarly, T cell depleted mice showed an approximate 94.62% decrease in total T cells compared to their IgG matched controls (Sup. Figure 3C, D).

**Supplementary Figure 3.**
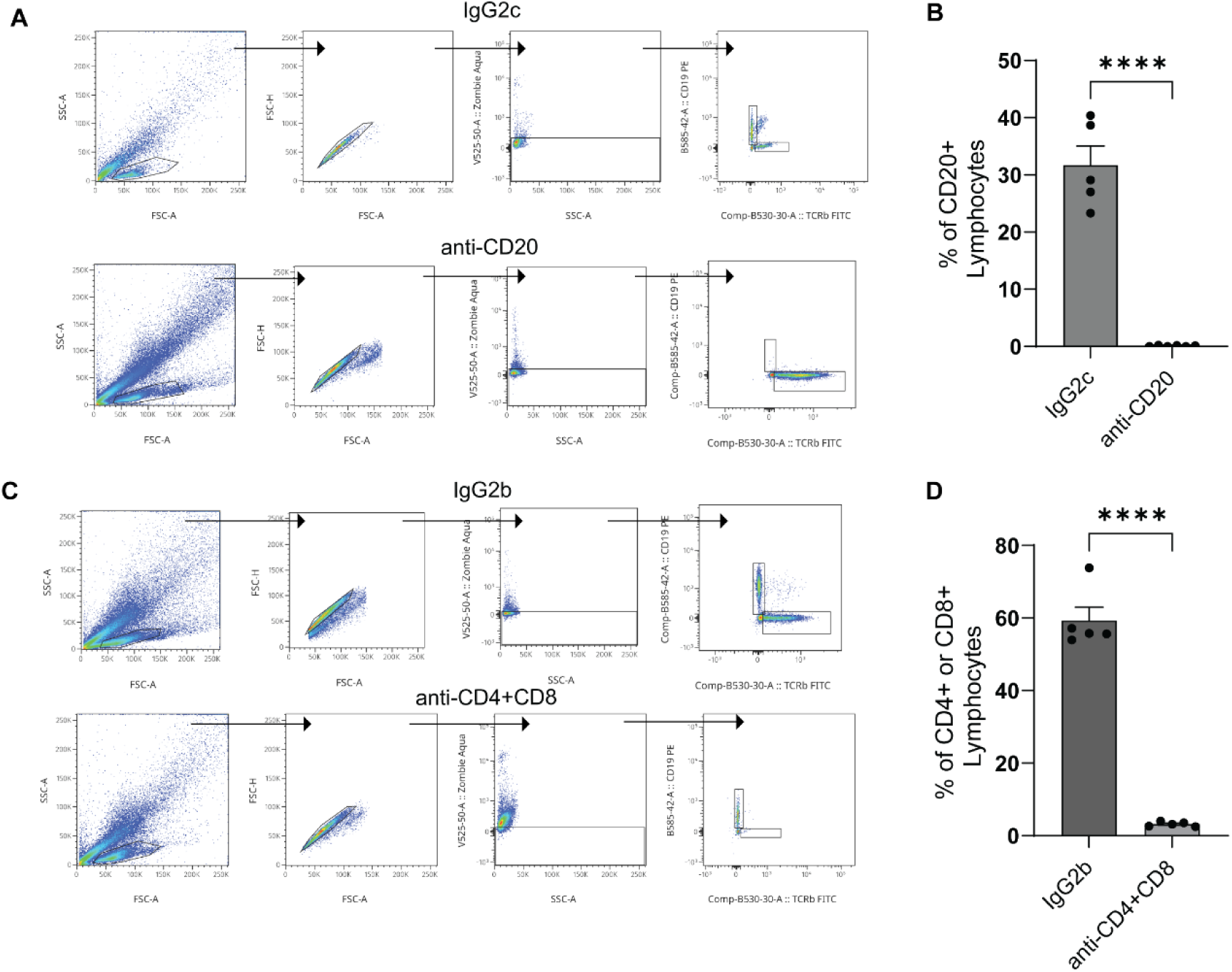
Treatment with *in vivo* CD20 or CD4 + CD8 antibodies successfully depletes B or T cell populations in mice. *A.* Gating for CD20 B cell populations in isotype control mice versus anti-CD20 treated mice. *B*. Quantification of B cell populations 14 d after retro-orbital injection of antibodies. *C*. Gating for CD3 T cell populations in isotype control mice versus anti-CD4 + anti-CD8 mice. *D*. Quantification of T cell populations 14 d after intraperitoneal injection of antibodies. n = 5. Each dot represents one animal. \**p* < 0.05; \*\**p* < 0.01; \*\*\**p* < 0.001; analyzed by unpaired t-test. ns = not significant. Error bars = ± SEM.

SN brain sections were processed for stereological quantification, where we found that T depleted mice showed increased number of DA neurons compared to their isotype control (Fig. 6B, C). Confirmation of TH^+^ cell death was evaluated via DAPI quantification, which showed an increase in DAPI^+^ cells in the T depleted group, though insignificant (Fig. 6D). Evaluation of TH^+^ cells co-labeled with p-α-syn revealed no significant difference between T IgG or T depleted mice (Fig. 6E,F). Prior to endpoint, mice had performed a series of behavioral assessments, as described previously. T depleted mice performed significantly better on the pole test compared to isotype controls, but no differences were found in either the grip strength or open field test (Fig. 6G-I). Considering that the T depleted mice had more DA neurons, but there were no significant differences in DAPI^+^ neurons, we cannot rule out that T cell depletion only protected DA levels in these neurons or other non-DA neurons. This could explain the skewing of the ratio of DA neurons to other subtypes, who were susceptible in the SN region.

Lastly, B depleted mice and their isotype controls were evaluated for DA death and p-α-syn inclusions in the SN, where we found no significant differences between the groups (Fig. 6J-M). Likewise, these mice showed no differences in capabilities among pole test, grip strength test, and open field testing (Fig. 6N-P). Together, these results suggest that T cells after injury modulate the susceptibility of DA neurons to neurodegeneration.

## Discussion

Here, we report that a one hit, closed-skull mTBI model causes detrimental outcomes for SN DA neurons, concomitant with an increase in peripheral immune cell infiltration. Our results in both Rag2 KO and T cell specific depletion models demonstrate that the loss of T cell infiltration and signaling within the brain parenchyma provides positive outcomes for this highly susceptible DA neuron population, and in turn, improves chronic behavioral outcomes. Transcriptional data from DA neurons 90 d after injury insinuates the potential interaction of T cell-associated IFN-γ with DA neurons to contribute to the observed cell loss and downstream deficits.

Data from our mTBI model recapitulates the findings from studies using more severe models of TBI, showing capability of producing a chronic degeneration of DA neurons [31, 32]. Our data from mice who were given mTBI prior to the seeding of PFFs suggests that most of the observed DA cell death is associated with the mTBI itself rather than triggering an earlier onset or heightened severity of degeneration associated with the PFFs. However, investigation of later time points would provide insight into whether a greater escalation of this loss occurs compared to mTBI or PFF only controls. With the goal of examining the impact of mTBI not only on DA neuron health and survivability, but also on α-syn propagation and seeding properties, we chose to generate PFFs from α- syn monomers. This model of rodent PD has been shown to produce LB pathology reflective of the human disease with DA neuron loss starting at 3 months post-injection [6, 33]. Previous TBI studies have involved models that overexpress endogenous α-syn. While these approaches are useful for producing robust aggregation and some symptomology, overexpression models often produce little to no neurodegeneration of midbrain DA neurons and artificial LB pathology, considered hallmark clinical features of PD [34, 35]. Similar studies have also employed MPTP or other pharmacological toxins to specifically ablate DA neurons. These models address the lack of degeneration; however, the highly accelerated speed of degeneration (hours/days) and lack of realistic LB pathology are not ideal for studying disease progression [36–38]. Hence, the mTBI and PD models chosen for our study reflect an improved translational perspective for how brain injury influences PD-associated pathologies and symptoms. Despite mTBI not increasing PFF-associated DA degeneration 60 d after intrastriatal injection, we did observe a greater spread of α-syn inclusions to other brain regions such as the ipsilateral cortex (Fig 3H), supporting the possibility of greater overall degeneration in mTBI + PFF animals at more chronic time points in other regions of the brain not evaluated in the scope of this study.

While our data in immunodeficient mice shows neuroprotection of DA neurons in the SN, we also observed a significant increase in α-syn inclusions both among the DA population and in TH^-^ cell populations, suggesting a complex interplay between the immune response and α-syn pathology. Due to the lack of T and B cell infiltration and surveillance in these mice, it is possible that, as a compensatory mechanism, microglia enter a more reactive but less efficient state, resulting in an increase of internalized and improperly cleared p-α-syn. It has been demonstrated that in the presence of T cells, microglia have reduced phagocytotic activity in a synucleinopathy model [39], supporting that dysregulation of this relationship could lead to such observations. In our depleted T-cell model, α-syn aggregate load was not significantly different between groups (Fig. 6E). However, we cannot rule out that other neurons preferentially succumbed in that region nor that the events that occur after acute injury – modulated by T cells – may affect aggregation, despite no observed DA neuronal deficits at 14 dpi.

Future work will be needed to tease out the specifics of the resident and peripheral immune interplay driving these mTBI-associated alterations in α-syn spreading, as recent data suggest microglia play an increasing role in T cell signaling during neurodegenerative disease. CD8^+^ T cells were shown to cause abnormal activation of microglia via IFN-γ in a mouse model of Alzheimer’s disease in which CD8+ T-cells also drove disease progression [40]. Genetic deletion of CD4^+^ T cells resulted in reduction of the CNS myeloid MHC-II response to α-syn in a rodent overexpressing model, consequently protecting DA neurons from cell loss associated with the model [14]. This was also seen in a model of tauopathy where microglia and T cell crosstalk dictated disease outcome [41]. Studies examining postmortem human brain tissue support an increase in T cell populations in PD. One such paper evaluated CD8^+^, CD4^+^, and CD3^+^ T cells in PD and associated parkinsonian disorders, finding that these patients show a range of increases in all three populations compared to healthy controls, and that the neuroinflammatory response appears to be increased earlier in the disease state and diminishes with greater neuron loss [42]. Further examination of microglia morphology and subsequent manipulation of the cell population, within the context of our experiments, could provide greater insight into whether they play a role in the observed deficits downstream of T cell signaling pathways as innate immune signaling pathways were also activated in our RiboDat dataset (Fig. 3).

Our data profiles the critical role of adaptive immune signaling in the pathophysiology of mTBI-mediated PD at chronic time points post-injury. We observed that T cells specifically play a detrimental role in DA neuron susceptibility and that mTBI drives heightened aggregate propagation/LB spread in the rodent PFF model of PD. Further elucidation of the effect that the innate immune timescale has on augmenting the adaptive immune response in the context of mTBI and PD will provide additional insight into the cellular and molecular mechanisms that compound risk in PD.

### Limitations of the Study

Future work will need to involve female animals to determine if the same adaptive immune mechanisms are at play, but considering that PD risk is biased towards males [43, 44], as well as the male bias in head injury [45, 46], this initial study was performed in males. Additional chronic time points evaluating DA neurodegeneration and spread of LB pathology are also warranted, where we anticipate that more differences may be found between behavior and DA neurodegeneration between groups. We also note that not all T cells are harmful, though we genetically and antibody depleted both main groups, with CD4^+^ T cells displaying neuroprotective properties in other contexts [47]. Additional work in understanding the T cell receptor repertoire and activity of subsets of T cells throughout injury is necessary, in addition to whether they differ in individuals with PD without injury, where, in PD, this work has begun to be explored [48].

## Materials and Methods

### Sex as a biological variable

Our study exclusively examined male mice because the disease modeled is more relevant in males, and male animals exhibited less variability in phenotype after brain injury. It is unknown whether the findings are relevant for female mice, but given recent findings in sex-specific neuroimmune responses, it is possible that female cohorts would yield different results [49].

### Animals

All mice were housed in a pathogen-free facility on a 12h light/dark cycle at Virginia Tech at a maximum of five per cage and provided standard rodent diet and water ad libitum. Experimental cohorts to be used for behavior studies were housed in a separate, reverse light/dark cycle room within the same facility. B6J.129(Cg)-Rpl22tm1.1Psam/SjJ (RiboDat), B6.Cg-Rag2tm1.1Cgn/J (Rag2 KO), and C57BL/6J WT mice were purchased from Jackson Laboratories. All mice were on C57BL/6J background. All mouse experiments were conducted in accordance with the NIH guide for the Care and Use of Laboratory Animals and under the approval of the Virginia Tech Institutional Animal Care and Use Committee (IACUC).

### Mild, Diffuse TBI Injury

Male mice aged 8-12 weeks were anesthetized with an intraperitoneal (IP) injection of ketamine (100 mg/kg) and xylazine (10 mg/kg) cocktail and then positioned in a stereotactic frame with ear bars. Body temperature was maintained at 37°C, monitored with both a rectal probe and a controlled heating pad. Nair was used to remove hair from the scalp, and any excess was removed thereafter with PBS to avoid irritation. Prior to using surgical scissors to expose the skull, the area was sterilized using three swabs of 70% ethanol followed by a betadine soak and a subsequent wash with 70% ethanol. A mild, diffuse head injury was induced directly on the skull at midline, directly between bregma and lambda, using an eCCI-6.3 device (Custom Design and Fabrication) equipped with a 5 mm flat impact tip. The injury was induced at 90° (flat impact to the skull), with a 5.0 m/s velocity, 1.5 mm impact depth, and 200 ms dwell period. The incision was closed with Vetbond tissue adhesive (3M), and the animals received a subcutaneous injection of Buprenorphine SR (1 mg/kg). The animals were then placed into a heated cage and monitored until they fully recovered from anesthesia. Sham animals received the same treatment and care, but without receiving the injury.

### Stereotactic Injections of PFFs

Full length WT mouse recombinant α-syn monomers were purchased from AlexoTech. Monomers were diluted in 1X PBS and assembly reactions were agitated in an Eppendorf orbital mixer (1,000 rpm at 37°C) for 5 d after which PFFs were harvested and stored at -80°C. Prior to injections, mouse α-syn PFFs were thawed and sonicated at room temperature using a probe sonicator (Fisher Scientific, Model Cl-18), amplitude: 30%, pulsing 1s ON 1s OFF for 1 minute (min). Mice were anesthetized via IP injection of ketamine (100 mg/kg) and xylazine (10 mg/kg) cocktail and situated within the stereotactic frame. Following sterilization and exposure of the skull, a craniotomy was performed in the right hemisphere to expose the injection site into the right dorsal STr (coordinates from bregma: AP: +0.2 mm; ML: -2.0 mm; DV: -2.6 mm from dura) at a rate of 0.2 µL/min using a 10µL Hamilton syringe. Mice were injected with 2.5 µL of PFFs (2 µg/µL) or 2.5 µL of PBS as a control, as described in the literature [33, 50, 51]. After the injection was complete, the syringe was left in place for 5 min before being slowly withdrawn.

### Adaptive Immune Cell Depletion

For B cell depletion, mice were injected with 250µg of CD20 antibody (BioXCell #BE0356) or IgG2c (Sydlabs #PA005171) control diluted in PBS via retro-orbital sinus once every 30 d, beginning 14 d after TBI or sham surgery. To deplete T cells, mice were injected with a cocktail of CD4 (BioXCell #BE0003-1) and CD8 (BioXCell #BE0061) antibodies (200ug each) or IgG2b control (Sydlabs #PA007144) diluted in PBS via IP injection 14 d after injury, followed by maintenance doses of 200ug of each antibody every 7 d.

### Levodopa Reversal

Mice were treated with l-3,4-dihydroxyphenylalanine (L-DOPA) (100 mg/kg) and carbidopa (12.5 mg/kg) (Sigma) administrated via IP injection 1 h before behavioral testing.

### Pole Test

Mice were placed facing upwards at the top of a metal pole (50 cm tall, 1 cm in diameter) completely wrapped in laboratory tape to provide grip. The mice were acclimated to the experiment environment and equipment with three trial runs the day prior to baseline recordings. Mice then performed the assessment at 30, 60, and 90 dpi. The total time that it took for them to turn, orient themselves downward, and descend the pole was recorded. Three trials were performed with each mouse, with the average of the second two trials being used for analyses. Times were limited to 90 seconds (s). Falling also constituted as a failure and counted as the maximum 90 s.

### Grip Test

Mice were acclimated to the experimental environment and equipment prior to data collection for 30 min. Briefly, mice were held by the tail and lowered towards a metal grid attached to the grip strength sensor (Bioseb). The sensor was set to peak mode, which enables the measurement of the maximal strength (force, in grams) exerted by the mouse. Once the mouse gripped onto the grid, the tail was gently pulled along the sensor axle until the grip was released. The value displayed on the sensor screen was recorded as the maximal grip strength. Each mouse performed the test in triplicate (5 m rest between tests), and the average of the results was divided by the individual mouse’s weight for normalization.

### Open Field Testing

Briefly, mice were acclimated to the testing room in their home cage with the lid open for 30 min. At the start of testing, mice were individually placed in the center of an opaque arena (43 cm x 43 cm x 43 cm) and recorded for 15 min by an overhead camera while they freely explored under red light. Video scoring was automatically calculated by the tracking software (EthoVision XT 17, Noldus), which provided measures for total distance, time moving/not moving, and time spent in specific, pre-determined regions of the arena (inner, inter, and outer).

### Immunohistochemistry

Mice were deeply anesthetized via IP injection of ketamine (500 mg/kg) and xylazine (10 mg/kg) and tissue was fixed by transcardial perfusion. Animals were perfused with ice-cold PBS with 20U/ml heparin followed by ice-cold 4% paraformaldehyde (pH 7.4) in PBS. Brains were then removed and stored overnight (ON) in 4% PFA at 4°C. The following day, brains were moved to 30% sucrose in 1X PBS at 4°C ON. Tissue was then frozen on dry ice and embedded in 30% sucrose in O.C.T (Tissue-Plus O.C.T Compound). Forebrain striatal tissue was sectioned at 30 µm thickness, while midbrain SN tissue was sectioned at 50 µm thickness through the entirety of the SN formation (-2.69 to -4.03 mm posterior to bregma) using a cryostat (CryoStar NX50, Thermo Scientific). Serial sections were collected 150 µm apart.

For immunohistochemistry, slides were incubated with 0.4% Triton X-100 in PBS for 30 min and then blocked with 5% BSA, 2.5% NGS for 1 h. Slides were then incubated at room temperature ON in primary antibody diluted 1:200 in block. The next day, slides were thoroughly washed with 1X PBS before being incubated for 1 h at room temperature in the appropriate fluorescent secondary antibody (ThermoScientific, JacksonImmuno) diluted 1:400 in PBS. Slides then went through an additional series of 1X PBS washes before being mounted with DAPI Fluoromount-G (Southern Biotech). Slides that required neuron staining were stained with Nissl (1:200 in block, NeuroTrace 640/660 Deep-Red Fluorescent Nissl, Invitrogen) prior to the mounting step. Representative images of both the STr and SN regions were taken at 10x on a Nikon C2 confocal microscope. Maximum intensity projections were created in ImageJ.

### Antibodies for Immunostaining

The following antibodies were used for this study: TH (Immunostar, cat #22941), p-α- syn S129 (abcam, ab51253), CD8 (abcam, ab217344), CD4 (abcam, ab183685), CD20 (abcam, ab64088), Nissl (Invitrogen, N21483).

### Optical Density

10x images of the STr stained with TH were loaded into ImageJ. Images were converted to 8-bit, and three evenly sized ROIs were randomly selected and measured. The three grayscale values were then averaged, with the result being the assigned optical density for that sample.

### Stereological Quantification

The StereoInvestigator software (MicroBrightField) was used by a blinded investigator to estimate the number of TH^+^, TH^+^/p-αsyn^+^, TH-/p-αsyn^+^, and DAPI^+^ cells within the SN.

Ten coronal serial sections spaced 150 µm apart and spanning the entirety of the SN were used for each animal (imaged at 40x using the fluorescent Teledyne Photometrics Kinetix high-speed sCMOS camera with a large image stitch). To approximate cell number, StereoInvestigator Optical Fractionator was used with a grid size of 150 µm by 150 µm and a counting frame of 100 µm by 100 µm. A contour was drawn around the SN for each serial section, and no less than 10 randomized counting frames were evaluated for each section. The estimated section thickness for 50 µm sections was approximately 30 µm due to post-staining shrinkage. The number of cells per contour/section, estimated section mounted thickness, section interval, and number of sections were all used to estimate the number of cells within the entire SN for each sample.

For striatal Nissl counts, one section of STr tissue for each animal around the same bregma was imaged at 10x using the Nikon C2 confocal. To approximate cell numbers, the Optical Fractionator was used with a grid size of 450 µm by 450 µm and a counting from size of 300 µm by 300 µm. A contour was drawn around the dorsal STr, using the corpus collosum as a reference for consistent imaging/contouring. The given parameters provided between 6-8 counting frames for each sample. Estimated number of cells were divided by volume to give a result of cells per mm^3^.

### High-Performance Liquid Chromatography

Neurotransmitter and metabolite quantification was performed by Vanderbilt University CMN/KC Neurochemistry Core Lab. Briefly, mice were sacrificed via PBS perfusion, the STr was harvested from freshly collected brains, and samples were subsequently frozen on dry ice before being stored at -80°C.

### Flow Cytometry

Mice were deeply anesthetized via IP injection of ketamine (500 mg/kg) and xylazine (10 mg/kg). 0.5 – 1mL of blood was drawn via cardiac puncture. Blood was incubated in 5 mL of ACK lysis buffer (BioVision) for 5 m to isolate white blood cells before being diluted with PBS and centrifuged at 300 x *g* for 5 m. Supernatant was discarded and the pellet was washed once in cold PBS before once again isolating the pellet. Cells were stained with 100 ul of zombie aqua (BioLegend) diluted in PBS at 1:100 for 20 min. Following incubation, cells were diluted with 1 mL of staining buffer (BioLegend) and spun down. After removing the supernatant, 500 ul of 4% PFA was added and gently mixed with the pellet before incubating at room temperature for 15 m. Following re-pelleting, cells were incubated in Fc block (Miltenyi Biotec) diluted in PBS 1:100 for 10 m. Lastly, cell pellets were again isolated and then stained with primary antibodies for 1 h, before being washed in cell staining buffer and transferred to flow tubes in PBS + 0.5% BSA to be analyzed using the BD FACSARIA Fusion Flow Cytometer. Primary antibodies used were as follows: TCRb FITC (1:200, Invitrogen #11-5961-81), CD19 PE (1:160, Biolegend #152407), CD8 PE-Dazzle594 (1:160, Biolegend #100761), CD4 PerCP-Cy5.5 (1:80, Biolegend #100433), CD49b APC (1:80, Biolegend #108909), and NKp46 BV421 (1:40, Biolegend #137612). Antibodies were diluted in cell staining buffer for primary incubation. FlowJo software was used for analysis.

### Ribotag Immunoprecipitation

Animals were deeply anesthetized with a ketamine (500 mg/kg)/xylazine (10 mg/kg) cocktail and hand perfused with cold PBS to remove blood. Lysate preparation and immunoprecipitation was performed as previously described [27]. Briefly, perfused mice were decapitated, and brains were removed for further dissection using a brain matrix to isolate STr tissue. Tissue was then homogenized using a Dounce tissue grinder filled with pre-chilled homogenization buffer. The lysate was then transferred to a microcentrifuge tube for a 10 min centrifugation at 10,000 x g, 4°C before the supernatant was collected. Immunoprecipitation was performed as described using anti-hemagglutinin (HA) antibody (BioLegend, cat #901513) and magnetic protein A/G beads (Pierce). Following a 4 h incubation at 4°C with antibody, lysate with antibody was added to the beads and incubated overnight at 4°C. The next day, samples underwent multiple washing and mixing steps with high-salt buffer before proceeding with RNA isolation as described in the Qiagen RNeasy kit, as per manufacturer’s instructions.

### RNA Sequencing

RiboDat IPs were collected as described above. For cortical tissue, animals were deeply anesthetized with a ketamine (500 mg/kg)/xylazine (10 mg/kg) cocktail and hand perfused with cold PBS to remove blood. Cortical tissue was rapidly dissected in ice-cold PBS and stored in RNAlater (Sigma-Aldrich) until RNA extraction. RNA was isolated with the RNeasy Mini Kit (Qiagen) per the manufacturer’s protocol. Isolated RNA was sent to MedGenome (MedGenome Inc.) for RNA sequencing using the Illumina TrueSeq stranded mRNA kit for library preparation and the NovaSeq system for sequencing. Cortex RNA-seq data has been deposited on GEO depository: GSE301952 and RiboDat striatum RNA-seq data has been deposited on GEO depository GSE297640.

Bases with quality scores less than 30 and adapters were trimmed from raw sequencing reads by Trim Galore (v0.6.4). After trimming, only reads with length greater than 30 bp were mapped to mm10 by STAR (v2.7.3a). Raw counts for each gene were output by HTSeq (v0.11.2). Raw counts were normalized and used to identify differentially expressed genes (DEGs) by DESeq2 (v1.42.1) in R Studio.

For RiboDat IP samples, genes were filtered by a p-value less than 0.05 and at least a 1.2-fold change to be considered differentially expressed. For cortical tissue samples, genes were filtered by an adjusted p-value less than 0.05 and at least a 1.5-fold change. All DEGs were used for GO enrichment analysis using R package clusterProfiler (v4.10.1) and org.Mm.eg.db (v3.18.0). The top 10 most significant biological processes (BP) terms were used to generate GO circle plots with R package GOplot (v1.0.2).

DEGs from RiboDat IPs, determined using the thresholds above, were submitted through Ingenuity Pathway Analysis (Qiagen) software for analyses of upstream regulators and top biological functions. The top 10 inhibited and top 10 activated upstream regulators were used to generate the associated bar plot. Top Biological Functions were determined and selected from the Diseases and Functions output, and visually reconstructed in BioRender.

### Real-time qPCR

cDNA was synthesized using iScript cDNA Synthesis Kit (Bio-Rad), and qPCR was run on the StepOnePlus System (Applied Biosystems) using TaqMan Fast Advanced Master Mix (Applied Biosystems), 10 ng of cDNA, and 0.5 µl of each 20x TaqMan Gene Expression Assay in a 10 µl reaction. Each sample was run in technical triplicates and expression levels were normalized to GAPDH. Fold change was calculated by comparative CT method. TaqMan Gene Expression Assays (ThermoFisher) are as follows: Mm00438388_m1; Slc6a3 (DAT), Mm00447557_m1; TH, Mm99999915_g1; GAPDH.

### Statistical Analyses

Data were analyzed with GraphPad Prism 10. Data were tested for normality using the Shapiro-Wilk test. Data that were normally distributed were analyzed by a student’s two-tailed t test for comparisons of two groups and a one-way or two-way ANOVA with Tukey’s or Sidak’s multiple-comparisons test to compare more than two groups, as appropriate. If data did not pass normality testing, non-parametric alternatives were applied (Kruskal-Wallis with Dunn’s test or one-sample Wilcoxon signed-rank test). Significance for non-parametric analyses are denoted by the pound symbol (#) instead of an asterisk. Differences were considered statistically significant at p < 0.05. Data are reported as mean ± SEM with n values given in the figure legend.

## Acknowledgements

We thank the Neurotrauma Consortium at Virginia Tech (AMP), R35GM142368 supplement (AMP) and CDMRP PD240020 (AMP) for supporting this work. We also thank the Flow Cytometry Core at Virginia Tech for their assistance.

## Author Contributions

CK planned and performed experiments, analyzed data, and wrote and revised the manuscript. JPM, BAL, SB, LH, LM, XW performed experiments. MLO analyzed data. MWB analyzed data. AMP conceived the project, planned experiments, analyzed data, secured funding for the project, and wrote and revised the manuscript. All authors read and approved the manuscript.

## Declaration of Interests

The authors declare no competing interests.

## Notes

### Competing Interest Statement

The authors have declared no competing interest.

